# Modeling of Epigenetic Modification-Induced Changes in CRX-dependent Genes *cis*-Regulatory Elements

**DOI:** 10.1101/179523

**Authors:** Reafa A. Hossain, Nicholas R. Dunham, Megan E. Harris, Taylor L. Hutchinson, Justin M. Kidd, Lindsay R. Kohler, Gregory J. Salamon, Alexander L. Schmidt, Madison D. Thomas, Raymond A. Enke, Christopher E. Berndsen

## Abstract

**Purpose:** DNA methylation is a well characterized epigenetic repressor of mRNA transcription in many plant and vertebrate systems. However, the mechanism of this repression is not fully understood. The process of synthesizing a strand of RNA from DNA, or transcription, is controlled by proteins that regulate RNA polymerase activity by binding to specific gene regulatory sequences. Cone-rod homeobox (CRX) is a well-characterized mammalian transcription factor that controls photoreceptor cell specific gene expression. While much is known about the functions and DNA binding specificity of CRX, less is known about how DNA methylation modulates CRX binding affinity to genomic cis-regulatory elements.

**Methods:** We used bisulfite pyrosequencing of human ocular tissues to measure DNA methylation levels of the regulatory regions of *RHO*, *PDE6B, PAX6*, and *LINE*. To describe the molecular mechanism of repression, we used molecular modeling to illustrate the effect of DNA methylation on human *RHO* regulatory sequences.

**Results:** In this study, we demonstrate an inverse correlation between DNA methylation in regulatory regions adjacent to the human *RHO* and *PDE6B* genes and their subsequent transcription in human ocular tissues. Docking of CRX to our DNA models shows that CRX interacts with the grooves of these sequences, suggesting changes in groove structure could regulate binding. Molecular dynamics simulations of the *RHO* promoter and enhancer regions show changes in the flexibility and groove width upon epigenetic modification. Models also demonstrate that changes to the local dynamics of CRX binding sites within *RHO* regulatory sequences which may account for the repression of CRX dependent transcription.

**Conclusion:** Collectively, these data demonstrate epigenetic regulation of CRX binding sites in human retinal tissue and provide insight into the mechanism of this mode of epigenetic regulation to be tested in future experiments.

## INTRODUCTION

Epigenetic modification of genomic DNA and associated histone proteins are crucial regulatory signals allowing eukaryotic cells the ability to adapt to dynamic environmental conditions^1^. DNA methylation is the covalent addition of a methyl group to the C-5 position of cytosine bases in genomic DNA. This addition is catalyzed by structurally distinct DNA methyltransferases (Dnmt) enzyme family members^2,3^. In plant and vertebrate genomes, DNA methylation is required for normal development and function of organisms^4,5^. DNA methylation has been linked to many key processes in vertebrate genomes, such as X-chromosome inactivation^6^, regulation of tissue-specific gene expression^7^, and suppression of mobile element transposition^8^. Dysregulation of DNA methylation-related epigenetic mechanisms is associated with human disease^9,10^. While DNA methylation has an increasingly appreciated role in complex genome regulation, the specific biochemical underpinnings of how this modification modulates the genome remain unclear.

Recent evidence demonstrates that DNA methylation regulates transcription within the retina. Cone and rod photoreceptor-specific genes display cell-specific patterns of DNA methylation, which appear to play an important role in the establishment and maintenance of retinal cell type-restricted gene expression^11,12^. Furthermore, targeted retina-specific disruption of Dnmts in murine models result in abnormal development of retinal neurons and dysregulation of global retinal gene expression^13–15^. Collectively these findings hint at an important role for epigenetic modification of DNA during retinal differentiation and maturation. However, deciphering the mechanistic detail of this role is vital for gaining insight into retinal function, retinal regeneration, and developing novel therapeutic strategies for retinal degeneration.

The homeodomain transcription factor cone-rod homeobox (CRX) is required for proper maturation of both rod and cone photoreceptors^16,17^. Mutations in CRX result in the blinding retinal degenerative diseases cone-rod dystrophy, Leber congenital amaurosis, and retinitis pigmentosa^18^. CRX mediates complex photoreceptor-specific transcriptional networks through physical interaction with evolutionarily conserved DNA sequence motifs^19,20^. Despite the presence of hundreds of thousands of these cis-regulatory elements in the mouse genome, less than 6,000 functional CRX binding regions (CBRs) have been identified in the murine retina. Furthermore, a subset of these CBRs display cell-type specific affinity in mouse rods and cones^19^. Beyond sequence context, genomic features responsible for regulating the spatial and temporal binding of CRX are poorly understood and represent a significant gap in our knowledge of photoreceptor development. One potential model for this differential binding is the dynamic epigenetic modification of the genome influencing local chromatin conformation and accessibility to transcriptional regulators such as CRX. Evidence in model systems demonstrates that several CRX-dependent genes have an inverse correlation between DNA methylation and gene expression^11,12^. Here, we expand on these findings by demonstrating a similar relationship for the first time in primary human retinal tissue. We also use molecular modeling to build evidence that reversible DNA methylation proximal to CRX binding motifs alters structural characteristics of the DNA double helix including minor and major groove width and DNA flexibility that may modulate CRX binding. Collectively, this study offers compelling evidence that DNA methylation plays a critical role in epigenetic regulation of human photoreceptor neurons and also provides insight into the biochemical interactions underlying this mode of regulation.

## METHODS

### Tissue Collection

De-identified post-mortem human donor eyes procured from three individuals with no reported ocular disease (National Disease Research Interchange, Philadelphia, PA) were used to collect ocular tissues for gene-specific quantification of DNA methylation. A scalpel was used to pierce the limbus followed by collection of the cornea using scissors. Whole corneas were flash frozen and ground into a fine powder using a mortar and pestle then immediately transferred to nucleic acid extraction buffer and stored at −80°C. Eyecups were further dissected with scissors by making four radial cuts exposing the retina. Cuts posterior to the ciliary margin were made liberating the retina from the anterior portion of the eye. Vitreous was removed and the retina was carefully peeled away from the eyecup using fine forceps. Whole retina from each donor was briefly washed in HBSS without calcium or magnesium to rinse away contaminating retinal pigment epithelial cells, placed in nucleic acid extraction buffer, vigorously vortexed, and stored at −80°C.

### Nucleic Acid Purification

Genomic DNA was extracted from human ocular tissues using a Qiagen Allprep Kit (Hilden, Germany) per manufacturer’s instructions. Briefly, lysates were homogenized using Qiashredder spin columns. Lysate flow through was then transferred to silica-based spin columns where genomic DNA and RNA were sequentially purified. Quality and quantity of DNAs were examined using UV spectrophotometry.

### Bisulfite Pyrosequencing

Quantitative analysis of DNA methylation was measured using bisulfite pyrosequencing performed as previously described^14,21^. Briefly, bisulfite conversion was performed on 200 ng genomic DNA using the EZ DNA Methylation-Gold Kit (Zymo, Irvine, CA). Following conversion, 30 μl PCR reactions were carried out using 2X JumpStart Taq Readymix (Sigma-Aldrich, St. Louis, MO). 5′ biotinylated PCR primers were designed to 5′ regulatory regions of target genes using PyroMark Assay Design 2.0 software (Qiagen, Hilden, Germany). Commercially available 5′ biotinylated PCR primers were used to amplify 5′ promoters of LINE1 retrotransposon repeats (Qiagen, Hilden, Germany). PCR cycling conditions were 95°C for 1 minute, followed by 45 cycles of 95°C for 30 seconds, 50-58°C for 30 seconds, and 72°C for 30 seconds, with a final extension at 72°C for 1 minute on a Bio-Rad C1000 Touch Thermal Cycler (Bio-Rad, Hercules, CA). Variable PCR annealing temperatures for different primer sets are indicated in Table 1. Biotinylated PCR products were purified and made single stranded to serve as a template in a pyrosequencing reaction using the PyroMark Q24 Vacuum Prep Tool per the manufacturer’s instructions (Qiagen, Hilden, Germany). A sequencing primer (0.3 μM final) was annealed to the purified single-stranded PCR product and pyrosequencing reactions were performed using the PyroMark Q24 Pyrosequencing System per manufacturer’s instructions (Qiagen, Hilden, Germany). Percent DNA methylation at each CpG dinucleotide in the BS PCR amplicon was determined and averaged between biological triplicates and statistical significance between the two sample groups was determined using a one-tailed t-test with a significance threshold set at 0.01. All PCR and sequencing primers used in these experiments are shown Table 1.

**Table 1.**
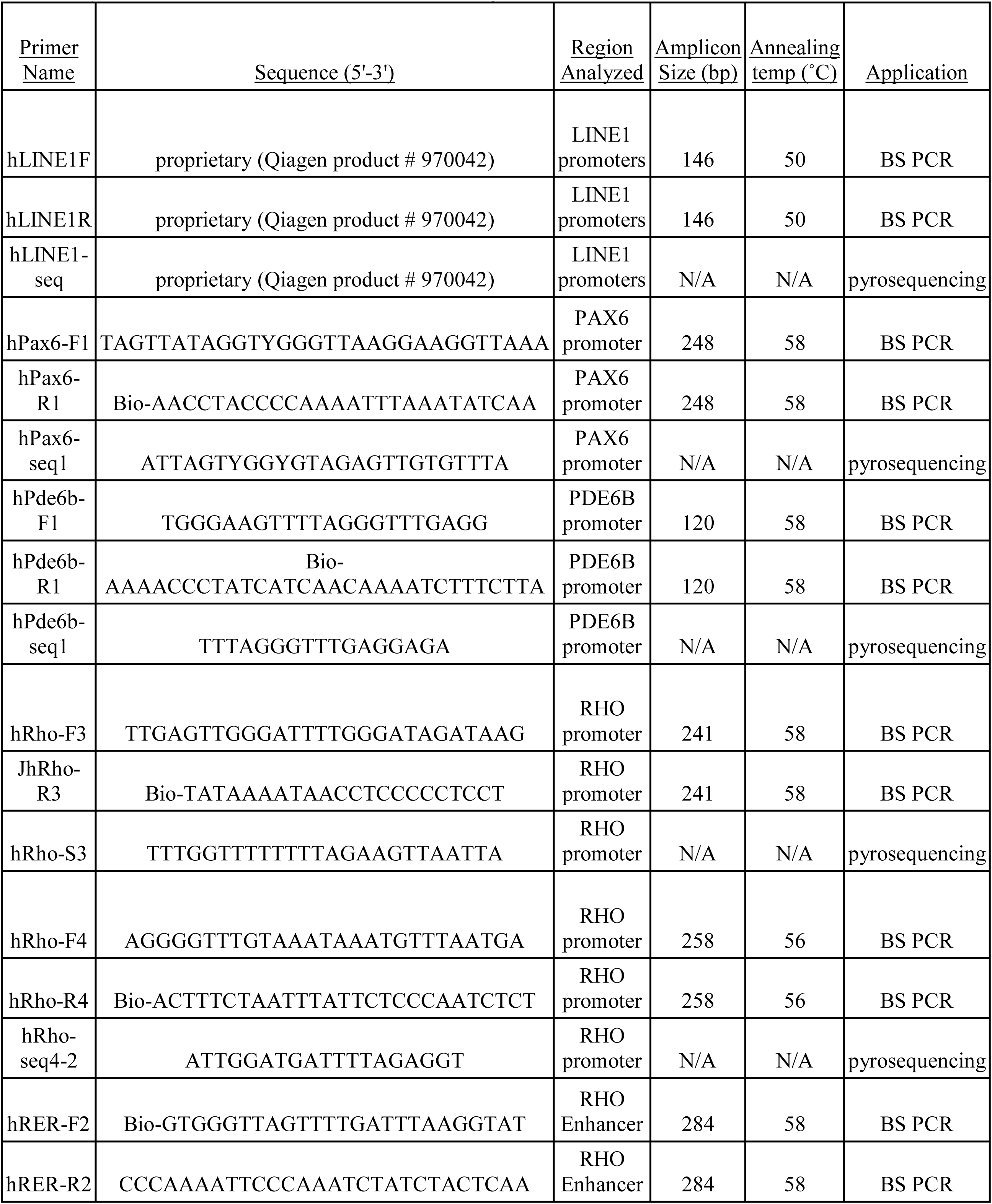
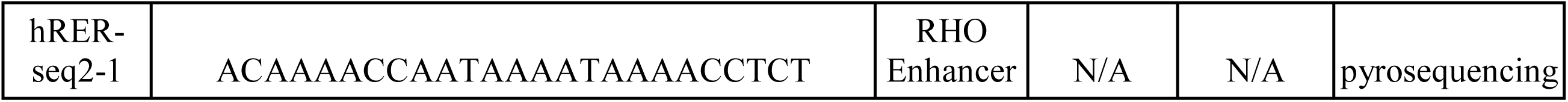
Oligonucleotides used for bisulfite PCR and pyrosequencing analysis. Bio indicates a biotinylation modification on the 5′ end of oligos.

### Bioinformatics Analysis of CRX Binding Sites

Computationally predicted CRX binding regions (CBRs) in the human genome were generated from CBRs obtained from a previously published study of mouse photoreceptors^19^. Experimentally validated mouse CBRs were aligned and mapped as a custom track to the human hg19 genome assembly using the LiftOver tool in the UCSC Genome Browser^22^. These data were overlaid with adult human retina RNA-sequencing transcriptome data as determined by Farkis et al., 2013^23^, vertebrate conservation data^24^, and custom tracks created in the UCSC Genome Browser highlighting various genomic sequences of interest^25^.

### Homology Modeling of Human CRX Binding Site

The UniProt accession number O43186 was used for modeling and assembly of CRX^26^. The DNA binding domain of CRX, consisting of amino acids 39-98, was generated using Swiss Model^27^. Models of the CRX DNA binding domains were energy minimized in YASARA before docking experiments^28,29^.

### Molecular Modeling of the Rhodopsin Enhancer Region and *RHO* Promoter Region

The RER enhancer region and *RHO* promoter sequences were obtained from UCSC Genome Browser and 3-D B-DNA structures generated using 3D-DART^30^ [REF-Genome Browser and 3D-DART]. The 39-bp DNA construct for the RER enhancer region consisted of the following bases: 5′-ACCTCATTAGCGTTGGGCATAATCACCAGGCCAAGCGCC-3′. The RHO promoter region consisted of the following 37-bps: 5′-TCTGCAGCGGGGATTAATATGATTATGAACACCCC-3′. DNA sequences were numbered in the 5′ to 3′ direction, continuing with the complementary strands. CpG methylation regions previously identified in this study were generated through YASARA. Methyl groups were built onto the C-5 position of the cytosine ring by using the “Swap>Atom” function. Molecular dynamics modeling was conducted using the AMBER14 force field in YASARA. The cut off for electrostatic interactions was set at 10.5 Å. The cell boundaries were periodic and filled with 0.3% magnesium chloride, and water molecules at pH 7.0. The simulation was run at 298.1 K. The save interval was every 0.1 ns over the 100 ns of simulation. All other parameters remained in the default setting.

RMSD, RMSF, and groove widths were calculated in YASARA. Groove widths were calculated by measuring the distance between the phosphate of one nucleotide to the phosphate of a nucleotide on opposite face of the groove. For the CRX binding sequence calculations, groove widths were localized to nucleotides 20-25 for the RER and 15-19 for the *RHO* promoter.

### CRX Docking

CRX was bound to RER and *RHO* sequences through HADDOCK to determine the interaction mechanism^31,32^. The previously modeled CRX DNA binding domain structure consisting of residues 39-98, was used in docking simulations with the energy minimized average structures from the promoter and RER simulations. Active residues for CRX include amino acids 40-46, 63, 69, 82, 84-85, and 88-93^26^. Active residues for the DNA molecules were bases 20-25 for the RER and 15-19 for the *RHO* promoter. CRX was then docked to the sequences using the Prediction interface on the HADDOCK server. We then used YASARA to determine the interactions occurring between the residues and the consensus sequence.

## RESULTS

### DNA methylation is inversely correlated with gene expression of photoreceptor-specific genes in human ocular tissues

Previous reports in the literature have demonstrated an inverse correlation between DNA methylation and gene expression of photoreceptor-specific genes^11,12,33^. These studies have been limited however, to murine model organisms or immortalized human cell lines and have yet to be characterized in primary human tissues. To determine if this trend is also observed in human ocular tissues, cornea and retina were collected from three sex-matched post mortem eyes procured from human donors 75 years or older (Figure 1). DNAs extracted from these tissues were used for quantitative bisulfite pyrosequencing analysis of DNA methylation on 5′ regulatory regions upstream of the phototransduction genes *RHO* (Figure 2A) and *PDE6B* (Figure 2B), the embryonic eye field transcription factor *PAX6*, and the multicopy long interspersed nuclear element (LINE1) repeats. The rod-specific genes *RHO* and *PDE6B* have been previously shown to have cell-specific patterns of gene expression in murine and human rod photoreceptors^11,12,34^. RNA-sequencing transcriptome data from Farkis et al., 2013 demonstrates that these rod-specific genes are highly transcribed in the adult human retina (Figure 2)^23^. Bisulfite pyrosequencing analysis of conserved 5′ regulatory regions upstream of *RHO* and *PDE6B* demonstrate lower levels of DNA methylation in retinal tissue relative to cornea, a tissue in which neither gene is expressed (Figure 3A+B). Levels of global DNA methylation between retina and cornea tissues were determined to be similar based on measurement of the constitutively methylated and silent LINE1 retrotransposon repeats as well as the unmethylated embryonic eye field transcription factor *PAX6* (Figure 3C+D). Collectively, these data demonstrate an inverse correlation between DNA methylation and transcription of *RHO* and *PDE6B* in human ocular tissues. These findings are consistent with previous observations of epigenetic regulation of phototransduction genes in the mouse retina as well as immortalized cell lines derived from human retinal tissue.

**Figure 1.**
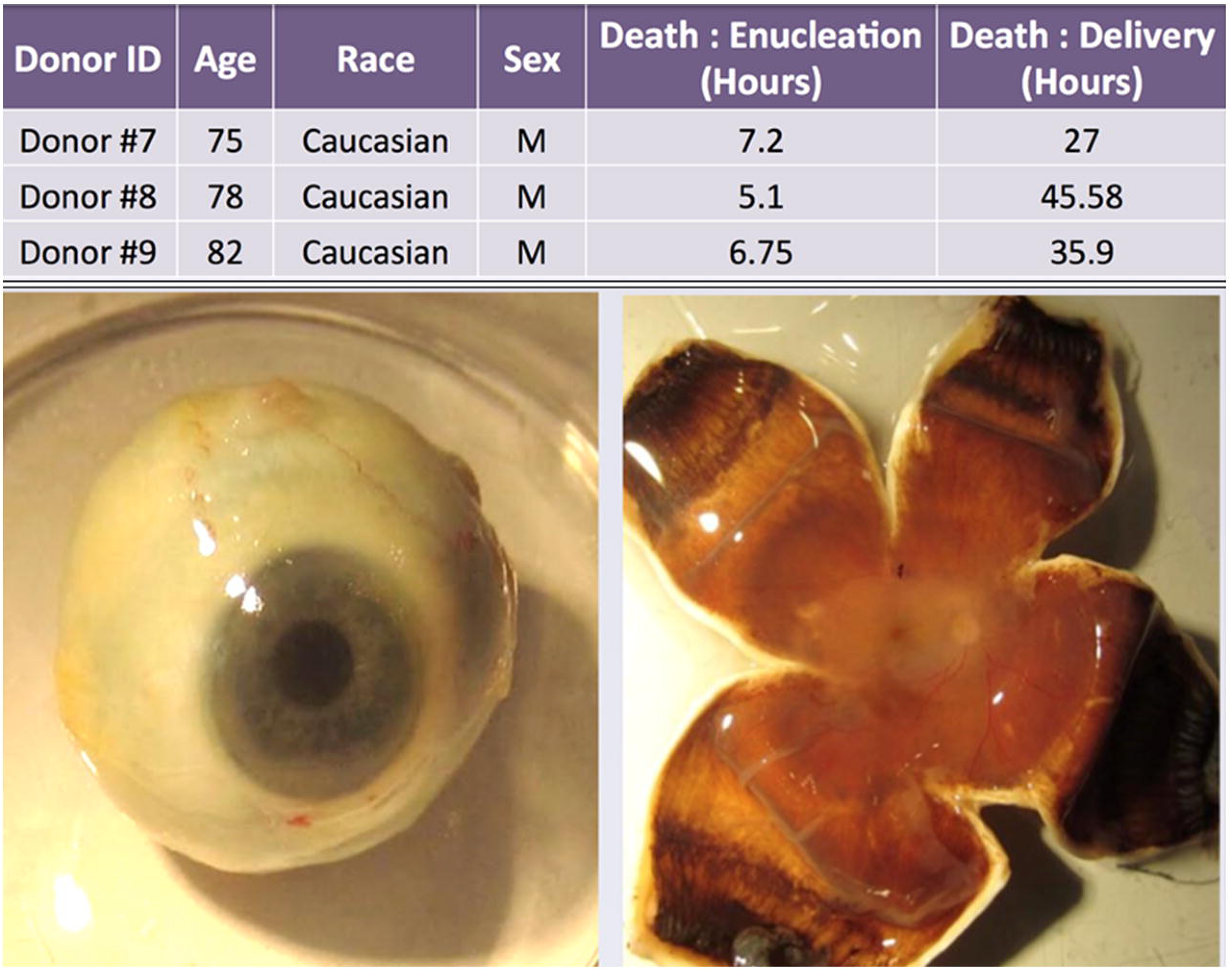
Donor eyes were excised within 10 hours from the time of death and delivered within 48 hours for the preservation of nucleic acids. Post-mortem human eye tissue collection strategy. **(left)** Whole globe image of human eye. **(right)** Cornea was dissected and collected followed by removal of anterior portion of the eye and flat mounting of the retina.

**Figure 2.**
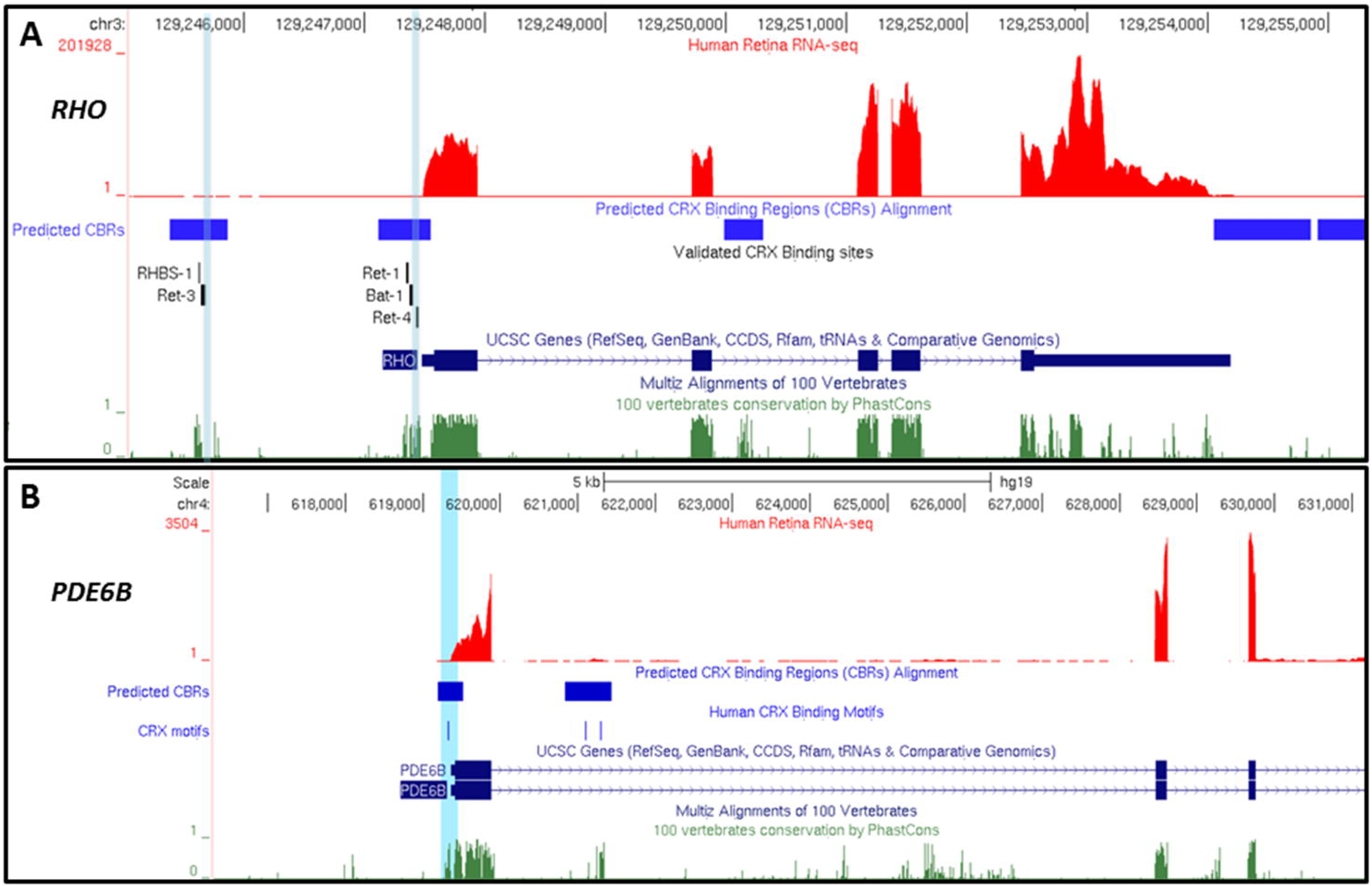
UCSC Genome Browser views of the *RHO* (A) and the 5′ region of *PDE6B* (B) photoreceptor-specific genes in the human hg19 genome assembly. Genes are oriented with the transcriptional start site on the left. From top to bottom, data tracks display 1) human hg19 genome coordinates, 2) human adult retina RNA sequencing data displayed as determined by Farkas et al,.2013^23^, 3) predicted CRX binding regions (CBRs), 4) CRX binding motifs present within predicted CBRs, 5) annotated genes and isoforms, and 6) evolutionarily conserved sequence averaged between 100 vertebrate species. Regions analyzed using bisulfite (BS) pyrosequencing are indicated with light blue highlights. An additional data track “Validated CRX/homeodomain Binding Sites” was added to the RHO locus representing sequences experimentally validated to bind to CRX and/or homeodomain family transcription factors^17,51^.

**Figure 3.**
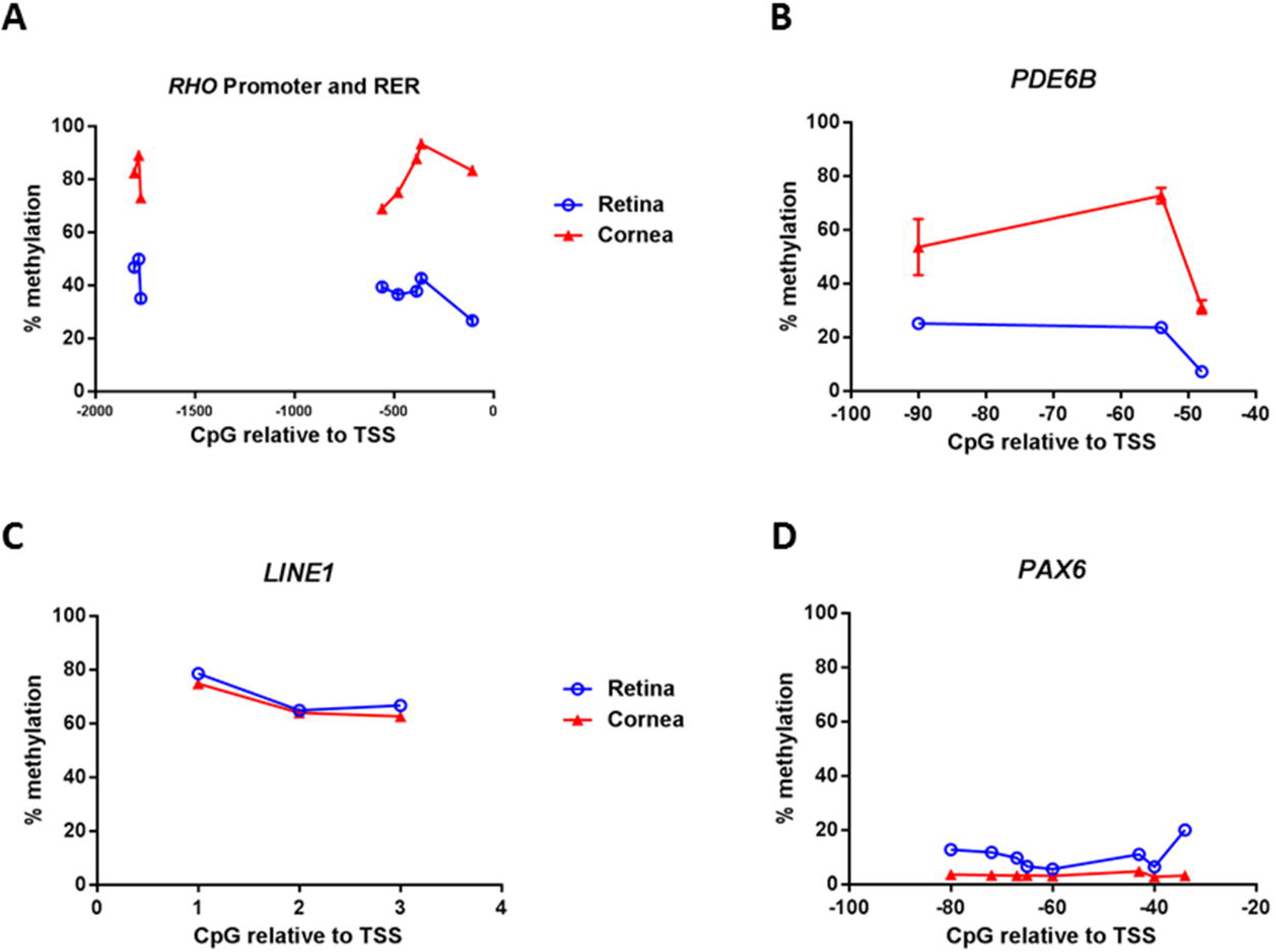
Quantitative bisulfite pyrosequencing analysis of DNA methylation at CpG sites relative to the transcriptional start site of **(A)** *RHO,* **(B)** *PDE6B*, **(C)** LINE1 repeats, and **(D)** *PAX6.* Data is presented as % Methylation at indicated genomic positions relative to the gene’s canonical transcriptional start site (TSS). Error bars represent standard error of the mean between three biological replicates of each sample (note that error bars are present but too small to see in panels A, C, and D. Statistical significance between retina and cornea at each CpG site was determined by t-test with a p<0.01. All CpG sites analyzed at all four loci were found to be have a p<0.01.

### Differentially methylated regions upstream of photoreceptor-specific genes correspond to CRX binding sites

Our epigenetic analysis of the human eye demonstrates tissue-specific patterns of DNA methylation in 5′ regulatory sequences of photoreceptor-specific genes that are inversely correlated with mRNA expression. These and previous observations in other mammalian retinal model systems suggests a functional role for DNA methylation in repressing transcription at these loci. However, a mechanism for this repression remains uncharacterized. A commonality between *RHO*, *PDE6B*, and many other photoreceptor-specific genes, is that they are transcriptionally regulated by the homeodomain transcription factor CRX^19,34,35^. This observation led us to question whether DNA methylation plays a role in regulating temporal and cell-specific binding of CRX to cis-regulatory regions within photoreceptor genomes. To test this hypothesis, we used a computational approach to align previously determined genome-wide CRX binding regions (CBRs) in the rod rich wild type mouse retina to the human hg19 2009 genome assembly (Figure 2; Predicted CBRs). To further assess the functionality of these presumptive regulatory regions, predicted CBRs were searched for experimentally validated CRX binding sites (Figure 2A; Validated CRX Binding Sites) or sequences containing CRX binding motifs (Figure 2B; Human CRX Binding Motifs). These analyses demonstrate that differentially methylated regions (DMRs) identified in this study are adjacent to experimentally validated CRX binding sites in the well-characterized *RHO* 5′ promoter and Rhodopsin enhancer region (RER), as well as predicted CRX binding sites in the *PDE6B* 5′ promoter. Given these results, we predict that differential methylation of these CpG sites may play a prominent role in modulating CRX binding to cis-regulatory elements upstream of *RHO*, *PDE6B,* and other CRX-regulated genes.

### The CRX DNA binding domain interacts with DNA grooves but not known methylation sites

DNA methylation is known to alter the three dimensional structure of double stranded DNA and interactions with DNA binding proteins such as transcription factors^36,37^. Computational modeling data suggest that the presence of a bulky methyl group in 5mC results in widening of the major groove and a concomitant narrowing of the minor groove^37^. Further, it has been experimentally validated that 5mC DNA bases biochemically mimic thymine bases due to similar hydrophobic interaction with major groove edge methyl groups^38^. More recent evidence has demonstrated a structural basis for affinity modulation of human transcription factors to methylated DNA via interactions of hydrophobic amino acid residues with methyl groups^36^. Taken together these observations suggest a mechanism for DNA methylation to narrow minor groove width and increase available hydrophobic interactions of cis-regulatory elements as a reversible mode of modulating CRX binding. However, a lack of structural information on CRX and the interactions of CRX with DNA hinders development of a specific mechanism. To test the hypothesis that methylated DNA sequences are less favorable binding partners with CRX than their unmethylated counterparts, we generated a homology model of the CRX DNA binding domain and models of the *RHO* promoter and the RER. The CRX binding domain model shows the expected 3 helix bundle characteristic of a helix-turn-helix homeobox protein (Figure 6). In both cases, DNA duplexes span one experimentally validated CRX binding site as well as one (promoter) or two (RER) CpG cytosine substrates for DNA methylation (Figure 4+5). We then docked the model of CRX to the DNA models using HADDOCK^32^. The CRX binding site within the promoter is within the minor groove and the C-terminal helix of CRX fits into the groove making contacts with the DNA backbone. Glutamic acid 80, lysine 88, and arginine 90 are known disease causing mutations within CRX and in our model of CRX bound to DNA, we see that K88 and R90 make electrostatic interactions with DNA (Figure 6)^39–41^. E80 makes contact with R69 and Q84, which contact the minor groove of DNA, suggesting a structural role for this amino acid (Figure 6). Thus, we felt confident that our CRX model could provide insight into how CRX interacts with DNA and is regulated by methylation. We next repeated the docking to the CRX binding sequence in the RER. The best interaction score produced by HADDOCK placed CRX in the major groove side of the CRX binding sequence. In addition to the previous backbone interactions noted for the promoter, CRX makes interactions with the DNA bases via N89 and K93 (Figure 6). Given that there currently is no structural information on CRX or these DNA sequences beyond the models we present here, we cannot exclude either binding mode. What is clear is that CRX likely interacts with a single DNA groove and does not make direct contact with methylation sites at this locus. These data suggest a mechanism of inhibition based on direct interaction with the methyl CpGs is unlikely.

**Figure 4.**
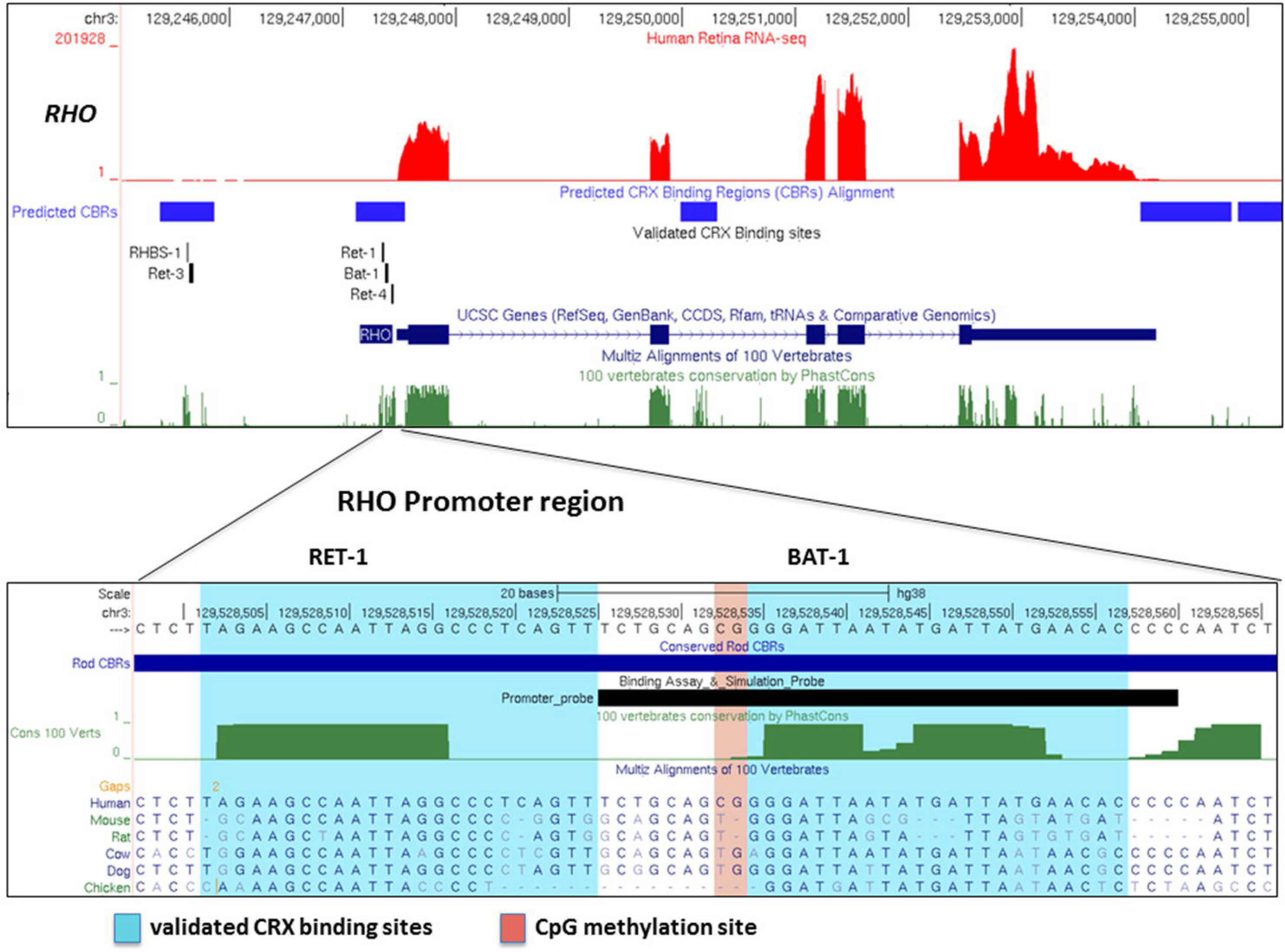
Top: UCSC Genome Browser views of the *RHO* locus in the human hg38 genome assembly oriented with the transcriptional start site on the left. From top to bottom, data tracks display 1) human adult retina RNA sequencing data displayed as determined by Farkas et al,.2013^23^, 2) predicted CRX binding regions (CBRs), 3) sequences experimentally validated to bind to CRX and/or homeodomain family transcription factors within predicted CBRs^17,51^, 4) annotated genes and 6) evolutionarily conserved sequence averaged between 100 vertebrate species. Bottom: Single base resolution view of the human RHO promoter aligned with five other vertebrate species. The experimentally validated CRX binding sites RET-1 and BAT-1^17^ are indicated with light blue highlights. The sequence used in CRX/promoter binding simulations is indicated as a black custom track.

**Figure 5.**
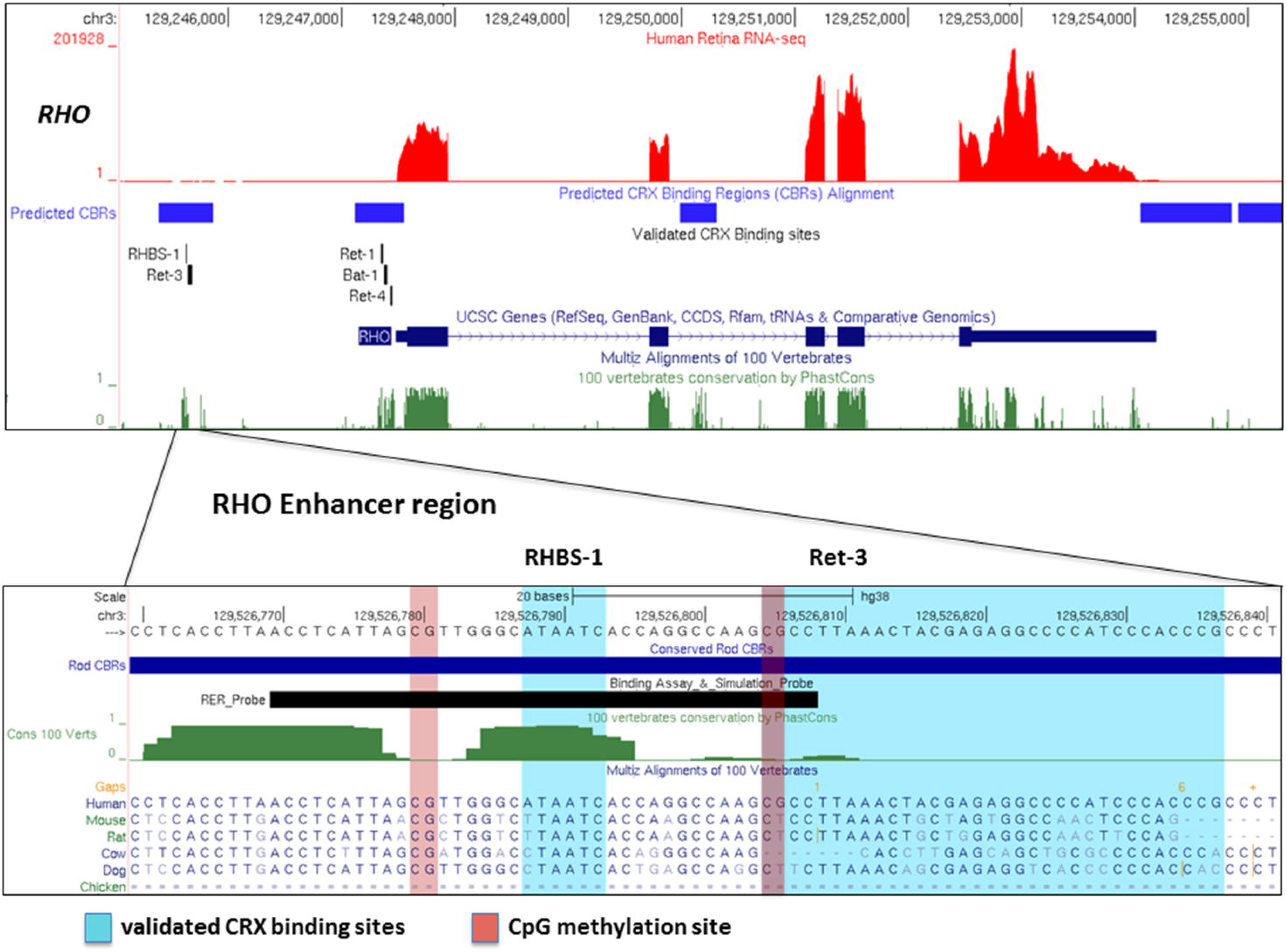
Top: UCSC Genome Browser views of the *RHO* locus in the human hg38 genome assembly oriented with the transcriptional start site on the left. From top to bottom, data tracks display 1) human adult retina RNA sequencing data displayed as determined by Farkas et al., 2013^23^, 2) predicted CRX binding regions (CBRs), 3) sequences experimentally validated to bind to CRX and/or homeodomain family transcription factors within predicted CBRs^17,51^, 4) annotated genes and 6) evolutionarily conserved sequence averaged between 100 vertebrate species. Bottom: Single base resolution view of the human RHO enhancer region aligned with five other vertebrate species. The experimentally validated CRX/homeodomain family transcription factor binding sites RHBS-1 and Ret-3^51^ are indicated with light blue highlights. The sequence used in CRX/enhancer binding simulations is indicated as a black custom track.

**Figure 6.**
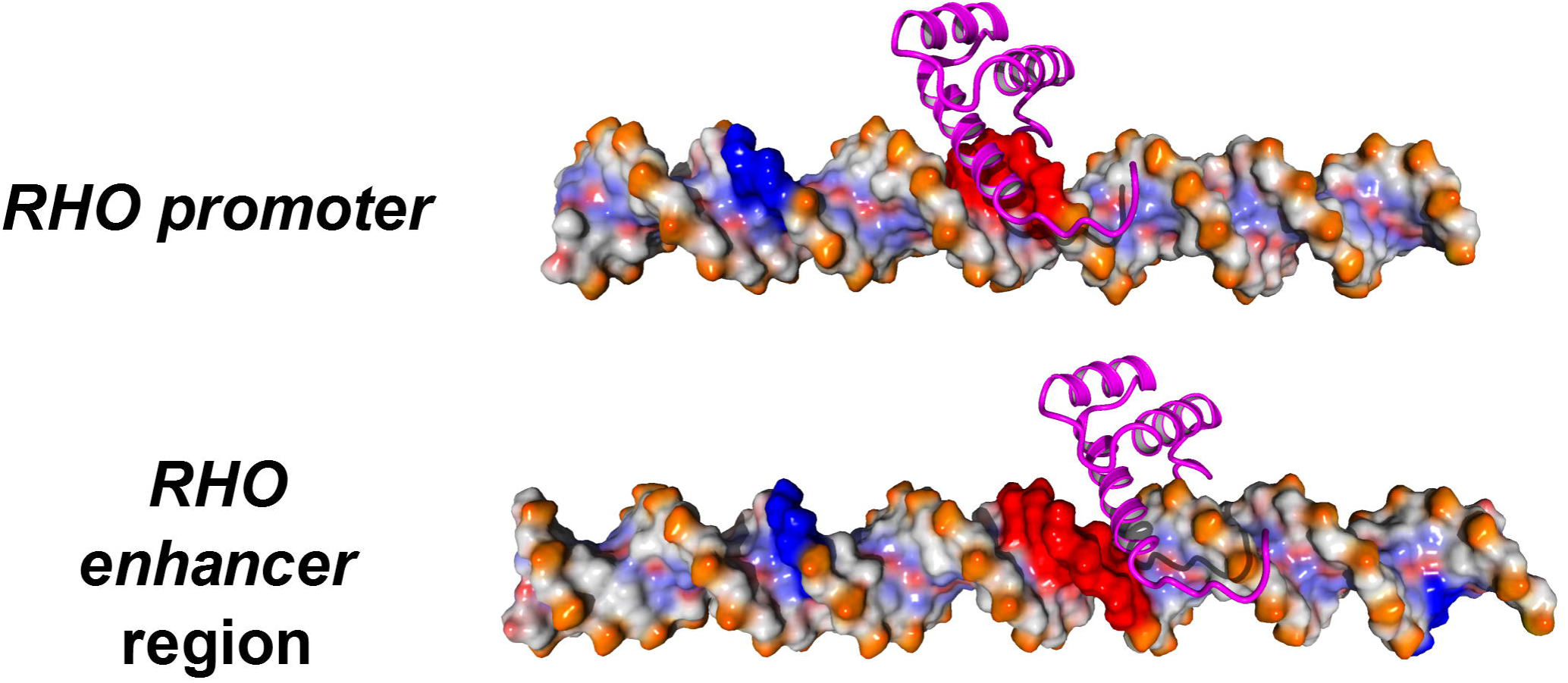
Model of CRX bound to the *RHO* promoter and RER. The CRX binding domain is shown in magenta, the methylation sites in blue and the CRX binding sequence in red.

### CpG Methylation enhances DNA flexibility

Our data showing that CpG methylation negatively regulates CRX binding raises the question of the molecular mechanism for binding inhibition. In addition to steric effects, CpG methylations have been proposed to change DNA dynamics and change the local structure of DNA, however which of these scenarios applies to CRX was not clear^36,37,42^. To address this question of the inhibition mechanism, we created a second set of DNA models that were methylated at the known sites and then equilibrated the models using molecular dynamics. We then analyzed the global and local effects of CpG methylation on the dynamics and structure of the DNA sequences and the CRX binding sites. Groove width statistics for these simulations are summarized in Table 2. While the mean and median values for the groove widths are similar, global analysis of the major and minor grooves shows that both structures are more dynamic (Table 2 and Figure 7). The increase in flexibility is indicated by the increased variance and standard deviations of the data measurements and visually by the wider distribution of values in the density plots (Figure 7 and Table 2). Analysis of the ensemble volume ratio shows that the structures of the methylated promoter have a larger volume, characteristic of higher dynamics. The effect of methylation is more apparent in the measurements that include just the CRX binding sequence, suggesting that the binding site is part of the dynamic region of DNA (Figure 7). These observations indicate that methylation does not alter the structure of DNA or the CRX binding site within the *RHO* promoter but changes the dynamics of the structure.

**Table 2.** Groove width statistics.

| Sequence | Modification | Groove | Mean width (Å) | Std. Dev (Å) | Median (Å) | Variance (Å) | Ensemble Volume Ratio<sup>a</sup> |
| --- | --- | --- | --- | --- | --- | --- | --- |
| RER | None | Major | 20.72 | 3.22 | 20.65 | 10.37 | 5.5 |
|  |  | Minor | 14.88 | 2.01 | 14.80 | 4.05 |  |
|  | Methyl | Major | 21.55 | 4.19 | 21.01 | 17.56 | 6.8 |
|  |  | Minor | 14.65 | 1.79 | 14.71 | 3.20 |  |
| Promoter | None | Major | 20.04 | 1.00 | 20.18 | 1.00 | 4.6 |
|  |  | Minor | 14.23 | 1.61 | 14.26 | 2.59 |  |
|  | Methyl | Major | 20.01 | 1.42 | 20.28 | 2.02 | 4.9 |
|  |  | Minor | 14.43 | 1.64 | 14.37 | 2.67 |  |

<sup>a</sup>Ratio of the combined volume of the ensemble of structures to the average volume of each structure in the ensemble

**Figure 7.**
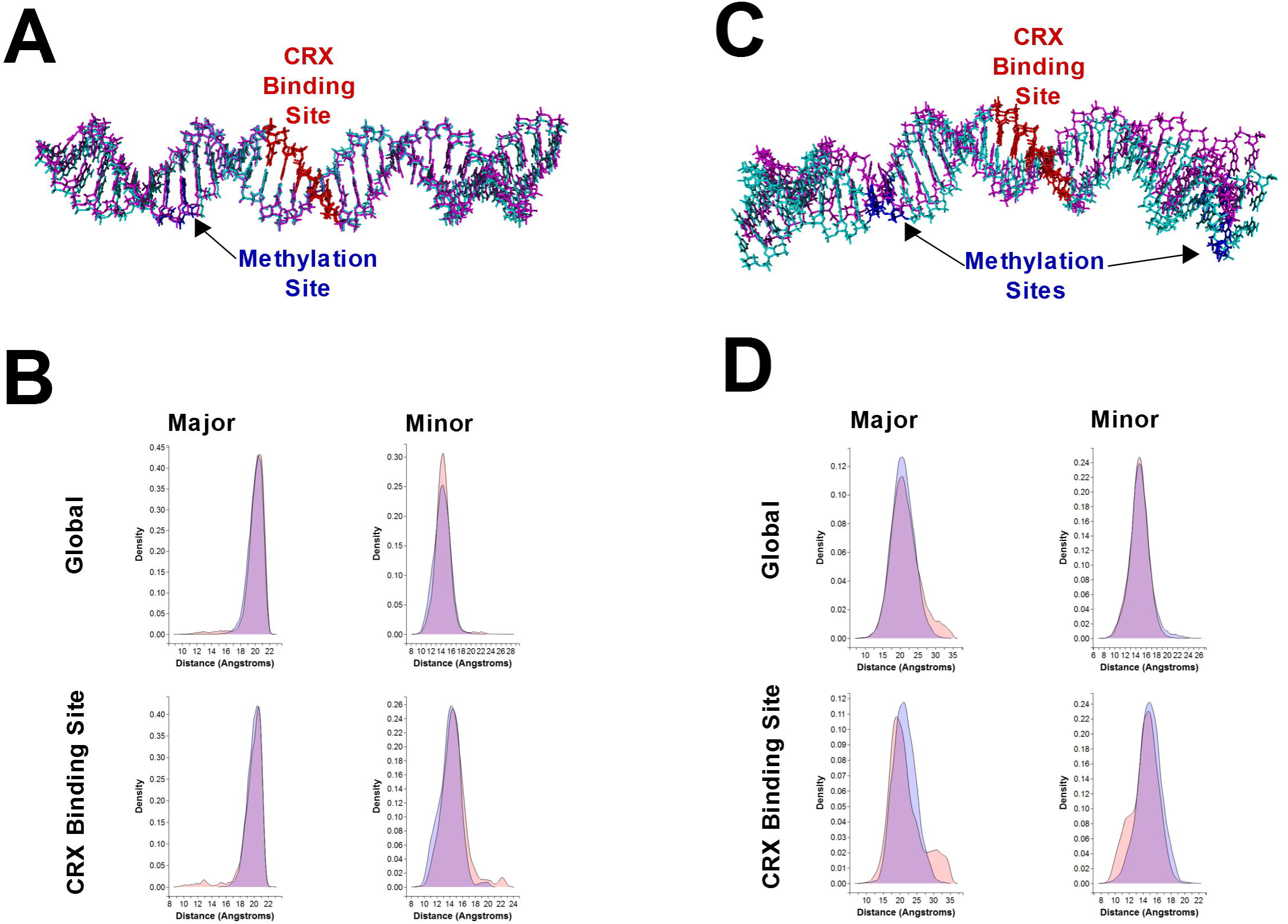
Structural and dynamic effects of methylation on the *RHO* promoter and RER **(A)** Alignment of unmodified (magenta) and methylated (cyan) *RHO* promoter DNA sequence. The CRX binding site is colored blue and the methylation site is colored red. **(B)** Analysis of the major (left) and minor (right) groove widths for the entire sequence (top) and the CRX binding site (bottom). Data from simulations of the methylated DNA are shown in red and data from the unmodified simulations shown in blue. Areas of purple indicate overlap. **(C)** Alignment of unmodified (magenta) and methylated (cyan) *RHO* enhancer region DNA sequence. The CRX binding site is colored blue and the methylation sites are colored red. **(D)** Analysis of the major (left) and minor (right) groove widths for the entire sequence (top) and the CRX binding site (bottom) Data from simulations of the methylated DNA are shown in red and data from the unmodified simulations shown in blue. Areas of purple indicate overlap.

We then performed these simulations and analyses using the RER sequence, which is methylated at sites on either side of the CRX binding motif. Methylation of the RER induced increased bending of the RER sequence in simulations relative to the unmodified sequence (Figure 7C). Further analysis of the groove widths in the RER sequence shows that with methylation the major groove widens while the minor groove narrows relative to the unmodified form (Figure 7 and Table 2). The standard deviation and variance for the major groove also increase, indicative of increased dynamics. The ensemble volume ratio increased by ~25% suggesting drastic changes in the dynamics of the RER upon methylation. The effect of methylation is especially apparent in the CRX binding site, which shows a significant population of major and minor groove widths in the methylated sequence simulations that is outside range values for the unmodified DNA sequence (Figure 7). Collectively, these data show that the presence of methylation alters the dynamics of DNA and the groove widths, possibly preventing efficient binding by CRX to the promoter and RER sequence.

## Discussion

Using cellular and *in silico* approaches, we addressed the regulation of CRX binding to genomic cis-regulatory elements and propose a model for how epigenetic DNA methylation may modulate this interaction. Bisulfite pyrosequencing of human ocular tissue showed that the *RHO* and *PDE6B* regulatory regions have lower levels of DNA methylation in the retina compared to corneal tissues (Figure 3). Given that CRX binds and regulates both genes, we postulated that CRX binding is inhibited by methylation of these regions upstream of the gene in non-expressing cells to block transcription. While this hypothesis will be best tested using biochemical methods, molecular modeling of the CRX DNA binding domain docked to CRX binding motifs within human *RHO* cis-regulatory elements were used in this study to demonstrate a basis for this model and to guide future biochemical experiments. To this end, our modeling data suggest that CRX interacts with the grooves of DNA and does not appear to make direct contact with methylation sites. Though CpG methyl sites do not occur within the CRX binding motifs analyzed in this study, molecular dynamics simulations demonstrated an increase in the overall structural dynamics and flexibility of CRX motifs adjacent to methyl CpG sites relative to data simulations of unmethylated sequences (Figure 7 and Table 2). Collectively, these data indicate that regional methylation of the *RHO* promoter and RER cis-regulatory elements may occlude CRX binding through alterations in the structure and dynamics of adjacent CRX binding motifs. An additional possibility unexplored in this study is that DNA methylation is an indirect effector of CRX affinity to cis-regulatory elements. Changes in DNA methylation are known to induce other repressive epigenetic modifications such as histone deacetylation and histone methylation^43,44^. Future studies focused on in vitro binding assays using methylated and unmodified oligonucleotides will be useful in testing a direct role of DNA methylation in CRX binding affinity.

The role of DNA methylation on CRX affinity has not been previously explored, however the effect of methylation on DNA structure and interactions with DNA binding proteins has been extensively studied. DNA methylation is known to affect local and regional DNA/nucleosome structure. Recent work indicates that DNA methylation both reduces DNA flexibility and enhanced nucleosome stability^42,45,46^. Confoundingly, other finding suggest that DNA methylation increases DNA flexibility or acts as a physical block to transcription factor binding^47–49^. In a broad sense, our results are consistent with methylation increasing DNA flexibility with a few important clarifications. The first clarification regards positioning of CpG methyl sites. Methylation sites investigated in this study are outside of the CRX binding motif and therefore likely do not act as a direct physical barrier to protein binding as characterized in other studies^36^. Alternatively, our findings indicate that CpG methyl sites adjacent to CRX binding motifs change the dynamics of the major and minor grooves over the entire sequence region including the CRX binding motif (Figures 7). This mode of epigenetic regulation may be used by cells to dynamically modulate binding of many different transcription factors at a particular regulatory locus and represents a potentially new paradigm for exploring the role of DNA methylation in regulating gene expression.

The second clarification regards motif-specific epigenetic modulation of CRX-DNA interactions. Previous studies have indicated that the specific mechanism of interaction between proteins and methyl CpGs appears to be highly sequence and factor dependent. Global analysis of the effects of DNA methylation on several transcription factors shows that CpG methylation can have positive, neutral, and negative effects suggesting that the molecular mechanism may be a combination of effects on specific proteins binding to specific loci^36^. Our data also support a model for DNA methylation influencing motif-specific effects on local DNA-protein interactions. Our molecular dynamics data indicates a methylated promoter CpG site is less perturbed than the effect observed for methylated RER CpG sites (Figures 7). Given that the *RHO* promoter and RER have little sequence similarity outside of conserved transcription factor binding motifs, it is not surprising that we observe distinct effects on structure and dynamics. In fact, recent functional evidence has demonstrated that individual CRX binding motifs within the murine *RHO* locus encode differential transcriptional responses to CRX ranging from strong inhibition of transcription to strong transcriptional activation^50^. Our data indicate that the methyl CpG adjacent to the CRX binding motif in the *RHO* promoter increases the width of the minor groove (Table 2). Inversely, methyl CpGs adjacent to the RER CRX motif increases the width of the major groove in our simulations. This discrepancy may be due to a number of sequence-related differences between the two cis-regulatory elements controlling human *RHO* transcription. Collectively, these data support the idea that the effect of DNA methylation on CRX binding is likely a combination of local and regional DNA sequence, distance of CpG methyl sites from CRX motifs, and the number of CpG methyl sites in or adjacent to CRX motifs. In this light, it is worth noting that though our data support a negative effect of DNA methylation on CRX binding at the *RHO* promoter and enhancer region, these finding do not preclude the possibility that DNA methylation may serve as a positive regulator of CRX binding at other loci in the genomes of retinal neurons. Future modeling studies of other CRX-regulated genes will be useful for further analyzing motif-specific epigenetic modulation of CRX affinity. Modeling data reported here as well as in future studies will guide precision biochemical analysis of the interaction between CRX with methyl CpGs at diverse photoreceptor-specific cis-regulatory elements.

## Acknowledgements

We wish to acknowledge C.E.B. as co-corresponding author who can be contacted at or R.A.E. can be contacted at. The authors would like to thank the students in CHEM361 for their efforts in the early stages of this project as well as Courtney Stout for her contributions to the project. This work was supported by NSF-REU (CHE1461175) awarded to the JMU Department of Chemistry and Biochemistry, Commonwealth Health Research Board grant #216-05-15 awarded to RAE, a JMU 4-VA Collaborative Research Grant awarded to RAE, and Burroughs Wellcome Fund Grant #1017506 awarded to RAE. Undergraduate students working on this project were also supported by endowed summer research scholarships provided by JMU alums Mr. and Mrs. Robert Crabtree and Jeffery E. Tickle to whom we thank for their generous support.

## References

1. Jenuwein T, Allis CD. Translating the histone code. Science (80-). 2001;293(5532):1074–1080. doi:10.1126/science.1063127.

2. Goll MG, Bestor TH. Eukaryotic Cytosine Methyltransferases. Annu Rev Biochem. 2005;74(1):481–514. doi:10.1146/annurev.biochem.74.010904.153721.

3. Cheng X, Blumenthal RM. Mammalian DNA Methyltransferases: A Structural Perspective. Structure. 2008;16(3):341–350. doi:10.1016/j.str.2008.01.004.

4. Vanyushin BF, Ashapkin V V. DNA methylation in higher plants: Past, present and future. Biochim Biophys Acta. 2011;1809(8):360–368. doi:10.1016/j.bbagrm.2011.04.006.

5. Jones PA, Takai D. The role of DNA methylation in mammalian epigenetics 8 227. Science (80-). 2001;293(0036-8075 (Print)):1068–1070. doi:10.1126/science.1063852.

6. Riggs AD. X chromosome inactivation, differentiation, and DNA methylation revisited, with a tribute to Susumu Ohno. Cytogenet Genome Res. 2002;99(1-4):17–24. doi:10.1159/000071569.

7. Goren A, Simchen G, Fibach E, Szabo PE, Tanimoto K, Chakalova L, Pfeifer GP, Fraser PJ, Engel JD, Cedar H. Fine tuning of globin gene expression by DNA methylation. PLoS One. 2006;1(1). doi:10.1371/journal.pone.0000046.

8. Slotkin RK, Martienssen R. Transposable elements and the epigenetic regulation of the genome. Nat Rev Genet. 2007;8(4):272–285. doi:10.1038/nrg2072.

9. Hansen KD, Timp W, Bravo HC, Sabunciyan S, Langmead B, Mcdonald OG, Wen B, Wu H, Liu Y, Diep D, Briem E, Zhang K, Irizarry RA, Feinberg AP. Increased methylation variation in epigenetic domains across cancer types. Nat Genet. 2012;43(8):768–775. doi:10.1038/ng.865.

10. Mikeska T, Craig JM. DNA methylation biomarkers: Cancer and beyond. Genes (Basel). 2014;5(3):821–864. doi:10.3390/genes5030821.

11. Merbs SL, Khan MA, Hackler L, Oliver VF, Wan J, Qian J, Zack DJ. Cell-specific DNA methylation patterns of retina-specific genes. PLoS One. 2012;7(3). doi:10.1371/journal.pone.0032602.

12. Montana CL, Kolesnikov A V, Shen SQ, Myers C a, Kefalov VJ, Corbo JC. Reprogramming of adult rod photoreceptors prevents retinal degeneration. Proc Natl Acad Sci U S A. 2013;110(5):1732–1737. doi:10.1073/pnas.1214387110.

13. Singh RK, Mallela RK, Hayes A, Dunham NR, Hedden ME, Enke RA, Fariss RN, Sternberg H, West MD, Nasonkin IO. Dnmt1, Dnmt3a and Dnmt3b cooperate in photoreceptor and outer plexiform layer development in the mammalian retina. Exp Eye Res. 2017;159:132–146. doi:10.1016/j.exer.2016.11.014.

14. Nasonkin IO, Merbs SL, Lazo K, Oliver VF, Brooks M, Patel K, Enke R a, Nellissery J, Jamrich M, Le YZ, Bharti K, Fariss RN, Rachel R a, Zack DJ, Rodriguez-Boulan EJ, Swaroop A. (Supp) Conditional knockdown of DNA methyltransferase 1 reveals a key role of retinal pigment epithelium integrity in photoreceptor outer segment morphogenesis. Development. 2013;140(6):1330–1341. doi:10.1242/dev.086603.

15. Rhee K-D, Yu J, Zhao CY, Fan G, Yang X-J. Dnmt1-dependent DNA methylation is essential for photoreceptor terminal differentiation and retinal neuron survival. Cell Death Dis. 2012;3:e427. doi:10.1038/cddis.2012.165.

16. Furukawa T, Morrow EM, Cepko CL, Ahmad I, Ahmad I, Redmond LJ, Barnstable CJ, Badiani P, Corbella P, Kioussis D, Marvel J, Weston K, Bally-Cuif L, Gulisano M, Broccoli V, Boncinelli E, Blitz IL, Cho KW, Bobola N, Hirsch E, Albini A, Altruda F, Noonan D, Ravazzolo R, Carter-Dawson LD, LaVail MM, Cepko CL, Austin CP, Yang X, Alexiades M, Ezzeddine D, Chen S, Zack DJ, Chiu MI, Nathans J, Chou W-H, Hall KJ, Wilson DB, Wideman CL, Townson SM, Chadwell LV, Britt SG, Conlon FL, Sedgwick SG, Weston KM, Smith JC, Dowling JE, Dryja TP, Li T, Fields-Berry SC, Halliday A, Cepko CL, Finkelstein R, Smouse D, Capaci TM, Spradling AC, Perrimon N, Fortini MK, Rubin GR, Freund CL, Gregory-Evans CY, Furukawa T, Papaioannou M, Looser J, Ploder L, Bellingham J, Ng D, Herbrick JS, Duncan A, al. et, Furukawa T, Kozak CA, Cepko CL, Hanes SD, Brent R, Hicks D, Barnstable C, Humphries MH, Rancourt D, Farrar GJ, Kenna P, Hazel M, Bush RA, Sieving PA, Sheils DM, McNally N, Creighton P, al. et, Jin Y, Hoskins R, Horvitz HR, Kikuchi T, Raju K, Breitman ML, Shinohara T, Kumar R, Chen S, Scheurer D, et al. Crx, a novel otx-like homeobox gene, shows photoreceptor-specific expression and regulates photoreceptor differentiation. Cell. 1997;91(4):531–541. doi:10.1016/S0092-8674(00)80439-0.

17. Chen S, Wang Q-L, Nie Z, Sun H, Lennon G, Copeland NG, Gilbert DJ, Jenkins NA, Zack DJ. Crx, a Novel Otx-like Paired-Homeodomain Protein, Binds to and Transactivates Photoreceptor Cell-Specific Genes. Neuron. 1997;19(5):1017–1030. doi:10.1016/S0896-6273(00)80394-3.

18. Rivolta C, Berson EL, Dryja TP. Dominant Leber congenital amaurosis, cone-rod degeneration, and retinitis pigmentosa caused by mutant versions of the transcription factor CRX. Hum Mutat. 2001;18(6):488–498. doi:10.1002/humu.1226.

19. Corbo JC, Lawrence KA, Karlstetter M, Res G, Corbo JC, Lawrence KA, Karlstetter M, Myers CA, Abdelaziz M, Dirkes W, Weigelt K, Seifert M, Benes V, Fritsche LG, Weber BHF, Langmann T. CRX ChIP-seq reveals the cis -regulatory architecture of mouse photoreceptors CRX ChIP-seq reveals the cis -regulatory architecture of mouse photoreceptors. 2010:1512–1525. doi:10.1101/gr.109405.110.

20. Lee J, Myers C a, Williams N, Abdelaziz M, Corbo JC. Quantitative fine-tuning of photoreceptor cis-regulatory elements through affinity modulation of transcription factor binding sites. Gene Ther. 2010;17(11):1390–1399. doi:10.1038/gt.2010.77.

21. Wahlin KJ, Enke RA, Fuller JA, Kalesnykas G, Zack DJ, Merbs SL. Epigenetics and cell death: DNA hypermethylation in programmed retinal cell death. PLoS One. 2013;8(11). doi:10.1371/journal.pone.0079140.

22. Speir ML, Zweig AS, Rosenbloom KR, Raney BJ, Paten B, Nejad P, Lee BT, Learned K, Karolchik D, Hinrichs AS, Heitner S, Harte RA, Haeussler M, Guruvadoo L, Fujita PA, Eisenhart C, Diekhans M, Clawson H, Casper J, Barber GP, Haussler D, Kuhn RM, Kent WJ. The UCSC Genome Browser database: 2016 update. Nucleic Acids Res. 2016;44(D1):D717–D725. doi:10.1093/nar/gkv1275.

23. Farkas MH, Grant GR, White JA, Sousa ME, Consugar MB, Pierce EA. Transcriptome analyses of the human retina identify unprecedented transcript diversity and 3.5 Mb of novel transcribed sequence via significant alternative splicing and novel genes. BMC Genomics. 2013;14(1):486. doi:10.1186/1471-2164-14-486.

24. Blanchette M, Kent WJ, Riemer C, Elnitski L, Smith AFA, Roskin KM, Baertsch R, Rosenbloom K, Clawson H, Green ED, Haussler D, Miller W. Aligning multiple genomic sequences with the threaded blockset aligner. Genome Res. 2004;14(4):708–715. doi:10.1101/gr.1933104.

25. Kent WJ, Sugnet CW, Furey TS, Roskin KM, Pringle TH, Zahler AM, Haussler D. The Human Genome Browser at UCSC. Genome Res. 2002;12(6):996–1006. doi:10.1101/gr.229102.

26. Bateman A, Martin MJ, O’Donovan C, Magrane M, Apweiler R, Alpi E, Antunes R, Arganiska J, Bely B, Bingley M, Bonilla C, Britto R, Bursteinas B, Chavali G, Cibrian-Uhalte E, Da Silva A, De Giorgi M, Dogan T, Fazzini F, Gane P, Castro LG, Garmiri P, Hatton-Ellis E, Hieta R, Huntley R, Legge D, Liu W, Luo J, Macdougall A, Mutowo P, Nightingale A, Orchard S, Pichler K, Poggioli D, Pundir S, Pureza L, Qi G, Rosanoff S, Saidi R, Sawford T, Shypitsyna A, Turner E, Volynkin V, Wardell T, Watkins X, Zellner H, Cowley A, Figueira L, Li W, McWilliam H, Lopez R, Xenarios I, Bougueleret L, Bridge A, Poux S, Redaschi N, Aimo L, Argoud-Puy G, Auchincloss A, Axelsen K, Bansal P, Baratin D, Blatter MC, Boeckmann B, Bolleman J, Boutet E, Breuza L, Casal- Casas C, De Castro E, Coudert E, Cuche B, Doche M, Dornevil D, Duvaud S, Estreicher A, Famiglietti L, Feuermann M, Gasteiger E, Gehant S, Gerritsen V, Gos A, Gruaz-Gumowski N, Hinz U, Hulo C, Jungo F, Keller G, Lara V, Lemercier P, Lieberherr D, Lombardot T, Martin X, Masson P, Morgat A, Neto T, Nouspikel N, Paesano S, Pedruzzi I, Pilbout S, et al. UniProt: A hub for protein information. Nucleic Acids Res. 2015;43(D1):D204–D212. doi:10.1093/nar/gku989.

27. Guex N, Peitsch M. SWISS-MODEL and the Swiss-PdbViewer: an environment for comparative protein modeling. Electrophoresis. 1997;18(15):2714.

28. Krieger E, Vriend G. New ways to boost molecular dynamics simulations. J Comput Chem. 2015;36(13):996–1007. doi:10.1002/jcc.23899.

29. Krieger E, Joo K, Lee J, Lee J, Raman S, Thompson J, Tyka M, Baker D, Karplus K. Improving physical realism, stereochemistry, and side-chain accuracy in homology modeling: Four approaches that performed well in CASP8. Proteins Struct Funct Bioinforma. 2009;77(SUPPL. 9):114–122. doi:10.1002/prot.22570.

30. van Dijk M, Bonvin amjj. 3D-DART: A DNA structure modelling server. Nucleic Acids Res. 2009;37(SUPPL. 2):235–239. doi:10.1093/nar/gkp287.

31. Van Dijk M, Visscher KM, Kastritis PL, Bonvin amjj. Solvated protein-DNA docking using HADDOCK. J Biomol NMR. 2013;56(1):51–63. doi:10.1007/s10858-013-9734-x.

32. Van Zundert GCP, Rodrigues jpglm, Trellet M, Schmitz C, Kastritis PL, Karaca E, Melquiond ASJ, Van Dijk M, De Vries SJ, Bonvin amjj. The HADDOCK2.2 Web Server: User-Friendly Integrative Modeling of Biomolecular Complexes. J Mol Biol. 2016;428(4):720–725. doi:10.1016/j.jmb.2015.09.014.

33. Mo A, Luo C, Davis FP, Mukamel EA, Henry GL, Nery JR, Urich MA, Picard S, Lister R, Eddy SR, Beer MA, Ecker JR, Nathans J. Epigenomic landscapes of retinal rods and cones. Elife. 2016; 5(MARCH2016). doi:10.7554/eLife.11613.

34. Peng G-H, Chen S. Active opsin loci adopt intrachromosomal loops that depend on the photoreceptor transcription factor network. Proc Natl Acad Sci U S A. 2011;108(43):17821–17826. doi:10.1073/pnas.1109209108.

35. Peng GH, Chen S. Crx activates opsin transcription by recruiting HAT-containing co-activators and promoting histone acetylation. Hum Mol Genet. 2007;16(20):2433–2452. doi:10.1093/hmg/ddm200.

36. Yin Y, Morgunova E, Jolma A, Kaasinen E, Sahu B, Khund-Sayeed S, Das PK, Kivioja T, Dave K, Zhong F, Nitta KR, Taipale M, Popov A, Ginno PA, Domcke S, Yan J, Schübeler D, Vinson C, Taipale J. Impact of cytosine methylation on DNA binding specificities of human transcription factors. Science (80-). 2017;356(6337):eaaj2239. doi:10.1126/science.aaj2239.

37. Machado ACD, Zhou T, Rao S, Goel P, Rastogi C, Lazarovici A, Bussemaker HJ, Rohs R. Evolving insights on how cytosine methylation affects protein-DNA binding. Brief Funct Genomics. 2015;14(1):61–73. doi:10.1093/bfgp/elu040.

38. Razin A, Riggs AD. DNA methylation and gene function. Science. 1980;210(4470):604–610. doi:10.1126/science.6254144.

39. Swaroop A, Wang QL, Wu W, Cook J, Coats C, Xu S, Chen S, Zack DJ, Sieving PA. Leber congenital amaurosis caused by a homozygous mutation (R90W) in the homeodomain of the retinal transcription factor CRX: Direct evidence for the involvement of CRX in the development of photoreceptor function. Hum Mol Genet. 1999;8(2):299–305. doi:10.1093/hmg/8.2.299.

40. Mitton KP, Swain PK, Chen S, Xu S, Zack DJ, Swaroop A. The leucine zipper of NRL interacts with the CRX homeodomain. A possible mechanism of transcriptional synergy in rhodopsin regulation. J Biol Chem. 2000;275(38):29794–29799. doi:10.1074/jbc.M003658200.

41. Nichols LL, Alur RP, Boobalan E, Sergeev Y V., Caruso RC, Stone EM, Swaroop A, Johnson MA, Brooks BP. Two novel CRX mutant proteins causing autosomal dominant leber congenital amaurosis interact differently with NRL. Hum Mutat. 2010;31(6). doi:10.1002/humu.21268.

42. Severin PMD, Zou X, Gaub HE, Schulten K. Cytosine methylation alters DNA mechanical properties. Nucleic Acids Res. 2011;39(20):8740–8751. doi:10.1093/nar/gkr578.

43. Du J, Johnson LM, Jacobsen SE, Patel DJ. DNA methylation pathways and their crosstalk with histone methylation. Nat Rev Mol Cell Biol. 2015;16(9):519–532. doi:10.1038/nrm4043.

44. Kondo Y. Epigenetic cross-talk between DNA methylation and histone modifications in human cancers. Yonsei Med J. 2009;50(4):455–463. doi:10.3349/ymj.2009.50.4.455.

45. Ngo TTM, Yoo J, Dai Q, Zhang Q, He C, Aksimentiev A, Ha T. Effects of cytosine modifications on DNA flexibility and nucleosome mechanical stability. Nat Commun. 2016;7:10813. doi:10.1038/ncomms10813.

46. Derreumaux S, Chaoui M, Tevanian G, Fermandjian S. Impact of CpG methylation on structure, dynamics and solvation of cAMP DNA responsive element. Nucleic Acids Res. 2001;29(11):2314–2326. doi:10.1093/nar/29.11.2314.

47. Geahigan KB, Meints GA, Hatcher ME, Orban J, Drobny GP. The dynamic impact of CpG methylation in DNA. Biochemistry. 2000;39(16):4939–4946. doi:10.1021/bi9917636.

48. Meints GA, Drobny GP. Dynamic impact of methylation at the M. HhaI target site: A solid-state deuterium NMR study. Biochemistry. 2001;40(41):12436–12443. doi:10.1021/bi0102555.

49. Carvalho ATP, Gouveia ML, Raju Kanna C, Wärmländer skts, Platts J, Kamerlin SCL. Theoretical modelling of epigenetically modified DNA sequences. F1000Research. 2015;4(0):52. doi:10.12688/f1000research.6148.1.

50. White MA, Kwasnieski JC, Myers CA, Shen SQ, Corbo JC, Cohen BA. A Simple Grammar Defines Activating and Repressing cis-Regulatory Elements in Photoreceptors. Cell Rep. 2016;17(5):1247–1254. doi:10.1016/j.celrep.2016.09.066.

51. Nie Z, Chen S, Kumar R, Zack DJ. RER, an evolutionarily conserved sequence upstream of the rhodopsin gene, has enhancer activity. J Biol Chem. 1996;271(5):2667–2675. doi:10.1074/jbc.271.5.2667.

